# Attraction of female house mice to male ultrasonic courtship vocalizations depends on their social experience and estrous stage

**DOI:** 10.1101/2023.04.27.538632

**Authors:** Jakob Beck, Bettina Wernisch, Teresa Klaus, Dustin J. Penn, Sarah M. Zala

## Abstract

Male house mice (*Mus musculus*) produce complex ultrasonic vocalizations (USVs), especially during courtship and mating. Playback experiments suggest that female attraction towards recordings of male USVs depends on their social experience, paternal exposure, and estrous stage. We conducted a playback experiment with wild-derived female house mice *(M. musculus musculus)* and compared their attraction to male USVs versus the same recording without USVs (background noise). We tested whether female attraction to USVs is influenced by the following factors: (1) social housing (two versus one female per cage); (2) neonatal paternal exposure (rearing females with versus without father); and (3) sexual receptivity (pro-estrous and estrous stages versus non-receptive metestrous and diestrous stages). We found that females showed a significant attraction to male USVs but only if females were housed with another female. Individually housed females showed the opposite response. We found no evidence that pre-weaning exposure to a father influenced females’ preferences, whereas sexual receptivity influenced females’ attraction to male USVs: non-receptive females showed preferences towards male USVs but receptive females did not. Finally, we found that individually housed females were more likely to be in sexually receptive estrous stages than those housed socially, and that attraction to male USVs was most pronounced amongst non-receptive females that were socially housed. Our findings indicate that the attraction of female mice to male USVs depends upon their social experience and estrous stage, though not paternal exposure. They contribute to the growing number of studies showing that social housing and estrous stage influence the behavior of house mice and we show how such unreported variables can contribute to the replication crisis.

## Introduction

House mice (*Mus musculus)* emit ultrasonic vocalizations (USVs) during courtship and mating, and the number of USVs emitted during sexual interactions correlates with males’ mating and reproductive success [1-2]. USVs have been described in wild-derived [3-4] as well as laboratory mice [5-9] in a variety of contexts. They are emitted as discrete sounds ("calls") that can be automatically detected [10] and classified into several different classes (3 to 12 "syllable" types), according to their shape, complexity, and other spectrographic features [11]. Male courtship USVs are uttered in repeated phrases that vary in syllable sequences ("syntax") and several other features comparable to the songs of birds and cetaceans [9]. Both sexes emit USVs during opposite-sex interactions [12] and their calls share similar qualitative features [13], though males produce most calls (ca. 85% to 93%) during opposite-sex interactions [14]. Male USVs are individually distinctive [15-16] and differ among laboratory strains [15]. Yet, males also modulate the emission of courtship USVs depending upon several factors, including their health [18], social housing [19], socio-sexual experience [20-22], sex of the stimulus individual [13], and estrous stage of a stimulus female [23]. Females’ responses towards male USVs have been investigated using playback experiments [2, 4, 24-29]. Females exhibit greater vocal responses to USVs than do males [26], and they are more attracted to complex than simple types of male USVs [25]. Females show more attraction to the USVs of males of their own than towards those of unfamiliar *Mus* species [27] and also towards USVs of unfamiliar non-kin versus familiar male siblings [4]. The aim of our present study was to investigate factors proposed to influence females’ attraction to male USVs.

The results of two studies have found three factors that appear to influence female attraction to male USVs (which interestingly are similar to factors proposed to influence female attraction to male scent [30-31]): Firstly, the habituation responses of female (CBA) mice to male USVs were influenced by whether they were kept in individual housing (IH) versus social housing (SH) [32]. Housing did not influence female attraction to USVs versus white noise, but IH mice and SH (with females) surprisingly showed increased attraction over time, whereas SH mice (with males and females) showed no increase over time. Social housing has been found to influence several behaviors in mice and other rodents [33], including sexual behaviors and auditory mechanisms that could alter female responses to USVs. For example, male mice [34] and rats (*Rattus norvegicus*) [35] kept in IH show lower sexual motivation than SH males, and female mice socially isolated during puberty exhibit less receptive sexual behaviors (lordosis) compared to socially reared individuals [36]. Mice kept in IH versus SH show differences in auditory perception [37], including perception of USVs [38], and neural auditory mechanisms [39-41]. These findings are interesting but also concerning because studies on laboratory rodents rarely provide information about social housing conditions, and such unreported variables potentially contribute to the replication crisis [42-45].

Secondly, another playback study found that female attraction to male USVs depended on whether females had been reared with their father (paternal exposure or +PE) or not (–PE) (C57BL/6 and Balb/C mice) [28]. Females were preferentially attracted to the USVs of males from strains that differed from their foster father, whereas females reared without a father lacked preferences. These results suggest that females learn characteristic features of their father’s USVs (sexual imprinting), potentially as a mechanism to avoid inbreeding [4]. This hypothesis predicts that neonatal paternal exposure will enhance female preferences for the USVs of unfamiliar males. These results also raise the possibility that neonatal exposure to the USVs of an adult male are necessary for females to develop a normal response to USVs (of any male). Studies on the effects of early social experience on behavior are needed, especially since PE and other rearing conditions are rarely reported in studies on laboratory rodents.

Thirdly, this same playback study found that females only showed preferences for male USVs when they were not sexually receptive (diestrous versus estrous stage) [28]. This result is surprising since females are expected to show enhanced attraction to male courtship signals, not a cessation of attraction during estrus. More studies are needed, however, as a previous study with wild house mice found estrus females were attracted to male USVs [4]. Estrous stage is another factor not usually reported in studies on laboratory rodents, and its effects on behavior are unclear and controversial [46].

We conducted playback experiments with wild-derived house mice (*Mus musculus musculus*) to examine the three factors proposed to influence female attraction to male USVs, which included: (1) social-versus individual-housing; (2) neonatal paternal exposure or not; and (3) sexually receptive (proestrus and estrus) versus non-receptive (metestrus and diestrus) stages. Females were simultaneously presented with a USV playback stimulus and a control (the same recording with USVs removed, leaving only the background noise). Trials were conducted in the absence of olfactory stimuli or direct male exposure, and level of females’ attraction to the two different acoustic stimuli was recorded. We expected that social housing (SH) and paternal exposure (+PE) would enhance female preferences for male USVs. Given that male USVs are courtship signals [3-9], we anticipated that sexually receptive females would show the strongest preferences, but that the opposite response was possible [28]. Finally, we recorded female habituation to male USV playbacks to determine whether the decreased interest in playbacks of male USVs over time found in most studies [4, 26, 28-29] depends on social housing [32].

We use the term “individual housing” rather than “social isolation,” because the mice were not socially isolated in the strict sense (as wthey were exposed to sensory cues from many other mice where they were kept in standard colony conditions). Also, we do not refer to social housing as the "control", as both types of housing are artificial treatments and neither are controls. IH and SH are both artificial conditions that lie in between two ends of a continuum of social conditions studied in rodents, from complete sensory and social isolation in artificial laboratory conditions on one end (social isolation *sensu stricto*) to social housing with both sexes in seminatural conditions at the other end (e.g., see Fig. S1 in [47]).

## Material and methods

### Animals

This study was approved by the Ethics and Animal welfare Committee of the University of Veterinary Medicine, Vienna in accordance with the University’s guidelines for Good Scientific Practice and authorized by the Austrian Federal Ministry of Education, Science and Research (BMBWF 2021-0.588.540) in accordance with current legislation. We conducted our study with 48 wild-derived female F2 generation mice as test subjects (aged 202-332 d). All mice were housed in standard colony conditions on a 12/12-hour light/dark cycle. Food and water were provided *ad libitum* with bi-weekly cage changes before vocal playbacks and individuals were earmarked for identification. All mice lived in standard mouse cages (Techniplast, mouse cage type IIL, 36.5 x 20 x 14 cm). Each cage contained aspen bedding (ABEDD, Austria), nesting material (Nestlet, Ehret, Austria), a translucent red house (Tecniplast, Germany) and a cardboard tube. At every cage change, 5 g of seeds and 5 g of apple were added to the cage as additional enrichment.

### Social experience treatments (rearing and housing)

We manipulated both paternal exposure and social housing, and we consider both treatments as types of social experience. Females were reared with their father (paternal exposure) or without their father, and weaning was conducted at 21 days after the birth of the pups. After weaning, juvenile mice lived in sibling groups for 14 more days and then female subjects were either socially housed (two female mice per cage) or individually housed (one mouse per cage). Female subjects belonged to one of four different treatments: 9 were individually housed at 5 weeks of age without their father present from birth until weaning (individually housed without paternal exposure, IH and –PE), 13 were individually housed at 5 weeks of age, but had their father present until weaning (individually housed with paternal exposure, IH and + PE), 13 were socially housed as adults but did not have their father present until weaning (socially housed without paternal exposure, SH and –PE), and 13 were socially housed as adults and had their father present until weaning (socially housed with paternal exposure, SH and +PE). Henceforth, we use abbreviations for the social treatment groups. Before weaning, mice lived with their litter mates, their mother, and the paternally exposed subjects with their father. Mice from all conditions lived in the same colony room and were exposed to visual, auditory, and olfactory signals from neighboring cages. None of the subjects had previous co-housing or other direct experience with males prior to the trials.

### Playback testing apparatus

Female preference assays were conducted using a Y-maze composed of three transparent plexiglass tubes connected by a non-transparent joint and the apparatus was lit from below with three infrared lights (CCD Camera, model DF-342A) (Fig. 1). The Y-maze was modified from simpler version designed for olfactory experiments [48] by connecting the stimulus arms of the maze to cylindrical arenas (covered with transparent covers on top and bottom), each having a hole for attaching speakers to present acoustic stimuli. Ultrasonic speakers (Ultrasonic Dynamic Speaker Vifa, Avisoft Bioacoustics, Germany) were protected from the mice with wire mesh and were positioned to project playbacks into the arenas and the arms of the maze. Equal loudness of the playback stimuli was confirmed by emitting the USV recording and control stimuli on both sides of the Y-maze, and measuring amplitudes (ultrasound microphone, Knowles FG, Avisoft Bioacoustics, Germany attached to an A/D converter, UltraSoundGate 116H, Avisoft Bioacoustics) at the joint of the arms prior to the trials. An ultrasound microphone was also placed next to the speakers to monitor playbacks from a laptop outside of the experimental room during trials.

**Fig 1.**
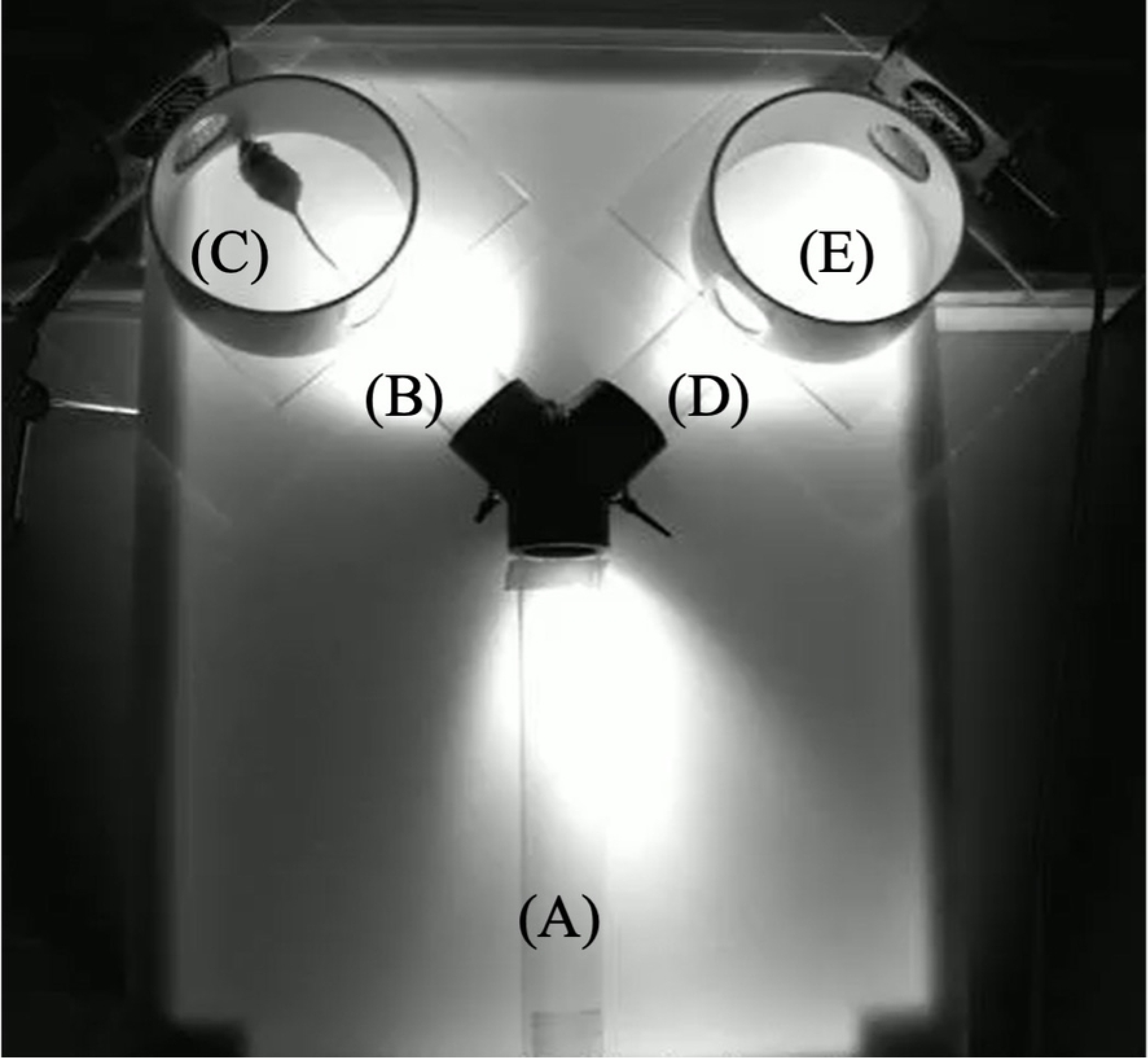
Y-maze apparatus for testing female preferences of playbacks of male USVs versus control. The maze was subdivided into the following sections: (A) neutral arm; (B and D) two stimulus arms and (C and E) circular arenas. Acoustic stimuli, either USVs or control (background noise) were broadcast into the arenas as well as the stimulus arms. For analyses, each stimulus arm and the connected circular arena were defined as one ’stimulus chamber’ (i.e., B and C versus D and E).

### Auditory stimuli: USV playback and controls

In each trial, a subject female was simultaneously presented with two different auditory stimuli: a recording of male USVs on one side of the maze and control "background noise" on the other side, i.e., the same audio file except with all USV syllables removed. This method ensured that the only differences in the acoustic stimuli were the male USVs. For USV playbacks, we used manipulated audio files created from recordings of male USVs and then played them back in a loop for five minutes into one of the two stimulus chambers of the Y-maze. We used 12 different recordings of 12 different males (F2 generation from a wild-derived population) that were unfamiliar and unrelated to the female subjects. Vocalizations of the males were recorded on two consecutive days under the following recording regime: On the first day, males were placed into a cage (type III, Tecniplast) with an unfamiliar and unrelated female from the same population that was divided by a perforated transparent plexiglass divider. Males were allowed to interact with the females through the divider for 10 minutes. Then, the plexiglass divider was removed, and males and females were allowed to interact directly for 10 minutes, as a method of sexual stimulation (priming) that elicits more vocalizations in the next recording [18]. In the final 5 minutes of recordings, the divider was returned, and males and females were returned to separate sections of the cage. All vocalizations were recorded using an ultrasound microphone (Knowles FG, Avisoft Bioacoustics, Germany). The same recording procedure was repeated per male on the next day, except a novel female stimulus was presented. The recordings that were utilized for the preparation of the playback files came from the first 10 minutes of recording on day 2 (i.e., primed males recorded for an unfamiliar female behind a divider). Recordings chosen to be used as playbacks were from the 12 males that emitted the highest number of USVs. We aimed to obtain files with >100 USVs to ensure feasible options for playback file modification. Of the 12 files used, 11 contained at least 116 USVs (range 116-130) but one with only 60 USVs was included to ensure that each of the four different housing treatment groups received the same playbacks. To further standardize playbacks, long pauses lacking male USVs within each recording were cut down to a pause length of 300 ms (previous recordings showed that the mean interbout duration was 300 ms in uncut male vocalizations), ensuring that subjects were presented with consistent auditory stimuli, without long pauses of silence between USVs. The final modified files were approximately 40 s long. Due to variation in the number of individuals within the female treatment groups, not all files were used once per treatment group. Females from the treatment group “IH and – PE” (n=9) received 9 unique playbacks, whilst the three other treatment groups (n=13) received the full 12 different playbacks with one vocalization being used twice. The creation of all USV playback and control playback files was conducted using the software STx (version 3.2, Acoustics Research Institute, Vienna). All sounds below 30kHz were removed from the final playback files using the software SASLab Pro (Avisoft Bioacoustics, Germany), thus excluding the frequencies that typically contain the most intense background noises, as well as audible vocalizations of mice.

### Experimental procedure

We provide a step-by-step protocol summarizing the necessary equipment and explaining how we conduct playback experiments (See S1 Methods in S1 File). Female subjects were sexually primed two days before the experiment with male bedding (15 g of a mixture of bedding from 10 males) and groups of 12 females were tested on eight different recording days. Each female was tested twice on two consecutive days, and all trials were conducted during the dark cycle under red light conditions. To begin each trial, subjects were gently transferred from their home cage to the Y-maze entrance (the neutral arm) using a handling bottle. Once the females voluntarily left the bottle and entered the Y-maze, a trap door was manually shut behind them to prevent returning to the bottle, and the experimenter left the room. The auditory stimuli (both USV and control) were already playing when the females were transferred into the Y-maze. On the second day of female testing, we switched the USV and background noise sides per female subject. To control possible side biases, the USV playback was also alternated between arms every four individuals so that half of the females were first tested with the USV stimulus on either side. Playbacks lasted a total duration of five minutes. Each mouse received its own Y-maze apparatus (clean tubes, lids, joints and arenas) to avoid contamination and the trap door was cleaned with ethanol after each trial. The body mass and a vaginal smear was taken from each subject immediately after removal from the Y-maze on trial day 2. Vaginal smears were evaluated using a light microscope and classified by their ovarian cycle as either sexual receptive (proestrus and estrus) or non-receptive (metestrus and diestrus). We tested the following female groups: IH and – PE (7 receptive, 2 non-receptive), IH and + PE (10 receptive, 3 non-receptive), SH and – PE (9 receptive, 4 non-receptive), SH and + PE (7 receptive, 6 non-receptive). Only data from the second trial day of the experiment were analyzed in the study, as this contained information about the body mass and receptive stage of the female, with the first trial serving as a way of habituating the females to the testing apparatus.

### Recording behavior

The subject females were video recorded during each trial using an infrared sensitive camera (D-Link, model DCS-3710) positioned above the Y-maze to capture all behaviors. Behavioral analyses were conducted using the Observer software (Observer XT 7.0, Noldus, Netherlands). Recordings were blindly analyzed by two experimenters at half speed. We define "chamber” as a stimulus arm and its corresponding circular arena. The duration of time spent in each zone of either side was recorded and summed to generate our response variables defined as “USV playback chamber” or “background noise chamber”.

### Statistical analyses

All statistical analyses were conducted using R (version 1.2.1335) [49] in the following steps: First, we tested whether females from the different treatment groups spent more time investigating the USV playback chamber or the control chamber and then we compared the intensity of their preference by comparing the ratio of time spent in the two different chambers across treatments (Side Preference Difference or SPD). SPD was measured as the time spent in the control chamber of the Y-maze subtracted from the time spent in the USV chamber:

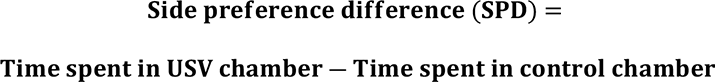

Thus, positive values indicate the female spent more time in the USV chamber of the Y maze, whilst negative values indicate they spent more time in the control chamber and the further the value from 0, the stronger the preference.

We compared the results for the entire five minutes of each trial and for the first minute, as previous studies have shown that females quickly habituate to playbacks [4, 26, 28-29]. Data for the one-minute analysis met the assumptions of normality and a linear mixed effect model (LMM) with a gaussian distribution was used to conduct a two-way mixed model Analysis of Variance (ANOVA) with random effects using the *CRAN* package *lmerTest (3.1-3).* Data over five-minutes did not meet the assumptions of normality and could not be transformed. Instead, a generalized mixed effect model (GLMM) was used with a gamma distribution taking the absolute value of time or SPD to conduct a two-way mixed model Analysis of Variance (ANOVA) with random effects using the *CRAN* package *lme4 (1.1-21)*.

To test whether females spent more time in either the USV or playback chamber, the fixed effect used in the LMM (over one minute) or GLMM (over five minutes) was chamber (USV playback and control), social experience (SH and + PE, SH and – PE, IH and + PE, IH and – PE) and receptivity (non-receptive or receptive) with mouse family and mouse ID as the random factors, and time as the response variable. Next, we looked at the effects of housing conditions (SH, IH) and paternal exposure (+ PE, – PE) as their own fixed effects along with receptivity (non-receptive or receptive), again with mouse family and mouse ID as the random factors. For this method, a significant interaction between chamber and one of the other fixed effects was needed to confirm a preference for either the USV or control chamber. To compare the intensity of their preference using SPD, a similar approach using both LMM (over one minute) and GLMM (over five minutes) was conducted. This time, SPD was the response variable, the same fixed effects were used, except the fixed effect “chamber” and the random effect “mouse ID” was removed from the model as using the response variable SPD rendered it redundant.

To compare female habituation to male playbacks, females from all different social treatments were analyzed separately to explore whether the effects of habituation differed across females from different social treatments. The ratio of the time spent in the USV playback chamber over the time spent in all three chambers (including neutral) was analyzed. To do this, a response variable named USV preference ratio (UPR) was calculated and defined as the total time spent in the USV chamber over the total time spent in all chambers.

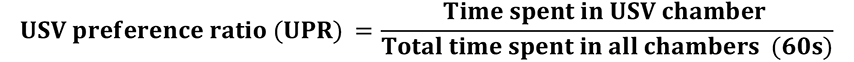

For testing, we used a LMM with UPR as the response variable, the fixed effect was minute block (1-5), and the random effect mouse ID was used to control for the repeated measures design of the analysis. Marginal R^2^ values were reported over conditional R^2^ values using the *r.sqauredGLMM* function from the *MuMin (version 1.43.17)* package, as the purpose of the analysis was to explore the fixed effect of minute block on UPR over any additional random effects. P values were obtained from summary result of the LMM.

We also explored whether housing conditions affected the body mass and estrous cycle of females and whether this would have any implications on female playback preference. Differences in body mass between the various social treatment groups were investigated using a LMM with the response variable body mass. The fixed effects chosen were either “overall social background”, “housing treatment” or “paternal exposure” and the random effect was age (to control for the variation in body mass not attributed to housing condition). To investigate whether the proportion of receptive or non-receptive females differed within housing treatments, binomial tests were conducted. To determine whether any differences in the proportion of receptive and non-receptive females were due to their housing treatment, we followed up binomial tests with a binomial LMM.

For all analysis, the choice of LMM or GLMM was subject to tests of normality and homogeneity of residual variance using residual and Q-Q plots as well as Shapiro-Wilk tests. LMM were preferentially chosen and data was transformed when necessary to do so. Only when data could not be transformed was GLMM preferred. For all models, we used a backwards stepwise model selection process and then subsequently compared their Akaike information criterion (AIC) to select for the model of best fit. On all occasions, interaction terms between the fixed effects were tested and then dropped if they were non-significant. If interaction terms were not significant but suggestive (p<0.08), we explored such interactions separately using a one-factor ANOVA design. In the present case, the effect of receptive stage on SPD was analyzed separately amongst IH and SH female treatment groups. All data were included across analyses and there were no outliers and results are show as means ± standard error (SE).

## Results

### Females’ overall preferences

First, we compared the differences in females’ responses to USVs among the four different social treatments (SH and + PE, SH and – PE, IH and + PE, IH and – PE). Our results overall indicated that the 48 females showed no significant preference towards for the USV playback or the control stimulus over the entire five min (F _(1,43)_ = 0.63, p= 0.432) or the first one min (F _(1,43)_ = 1.32, p= 0.256). There was also no significant interaction between the females’ overall social housing and their preference for either the USV playback chamber or the noise chamber at five min (F _(3,43)_ = 1.12, p= 0.295) or at one min (F _(3,43)_ = 1.64, p=0.194). There was a significant difference in SPD from females of different social treatments over five min (F _(3,44)_ = 2.95, p= 0.043), although Tukey post hoc tests revealed no pairwise contrasts at α <0.05. This result was not significant and consistent with the first min (F _(3,40)_ = 2.21, p= 0.102).

Next, we investigated whether females habituated to male USV playbacks. Collectively, females spent less time in the USV chamber over time (R_2_ = 0.06, p<0.001). This pattern was observed across all treatment groups; SH and + PE (R^2^ = 0.04, p=0.026), SH and – PE (R^2^ = 0.07, p<0.01), IH and + PE (R^2^ = 0.04, p=0.026), IH and – PE (R^2^ = 0.07, p<0.01).

There were no significant differences in body mass between females from the four different overall social housing treatments (IH and – PE, IH and + PE, SH and – PE, SH and +PE), (F _(3,42)_ = 1.09, p = 0.362).

### Social housing treatments

There was no significant interaction between housing condition and preference for either the USV playback or control chamber over five min (F _(1,46)_ = 2.86, p= 0.098) although there was a significant interaction over the first min (F _(1,46)_ = 5.25, p= 0.027). SH females spent more time in the USV playback chamber than IH females. SH females had a significantly higher SPD than IH females over both five min (F _(1,44)_ = 5.85, p= 0.020) and over the first minute (F _(1,43)_ = 6.64, p= 0.015).

There was no significant difference in body mass between IH and SH females (F _(1,45)_ = 3.12, p= 0.084).

### Neonatal paternal exposure

There was no significant interaction between paternal exposure and female preference for the USV playback stimulus or the control stimulus over five min (F _(1,44)_ = 0.02, p=0.915), or during the first min (F _(1,44)_ = 0.88, p= 0.767). There was no significant difference in SPD between females from different paternal exposures over five (F _(1,44)_ <0.01, p=0.980) or one min (F _(1,43)_ = 0.16, p=0.691). There was no significant difference in body mass between + PE and – PE females (F _(1,44)_ = 0.07, p= 0.800).

**Figure 2.**
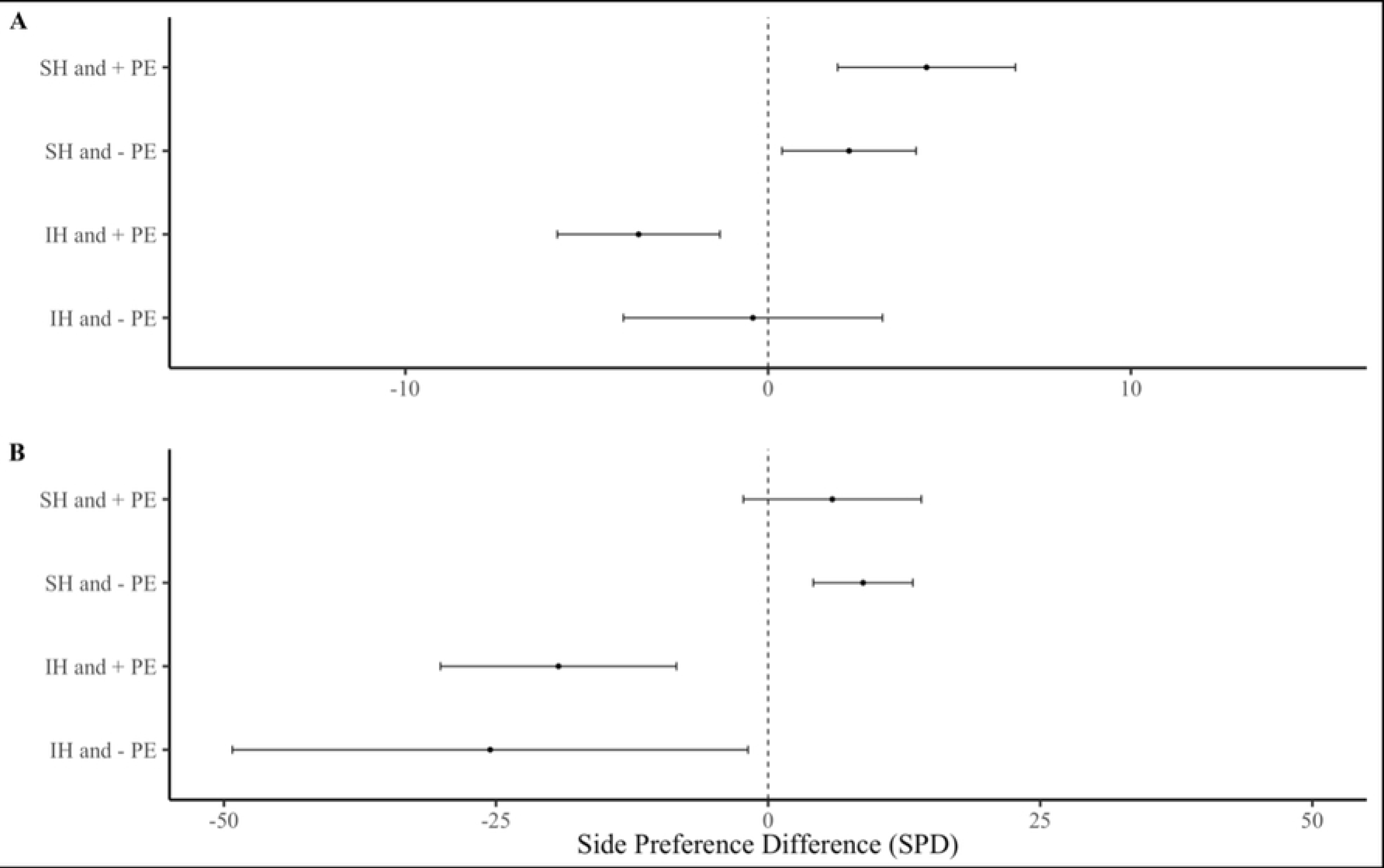
Side preference difference (SPD) of females from different social treatments. Showing **(A)** over one min and **(B)** over five mins. Error bars show mean ± SE.

**Figure 3.**
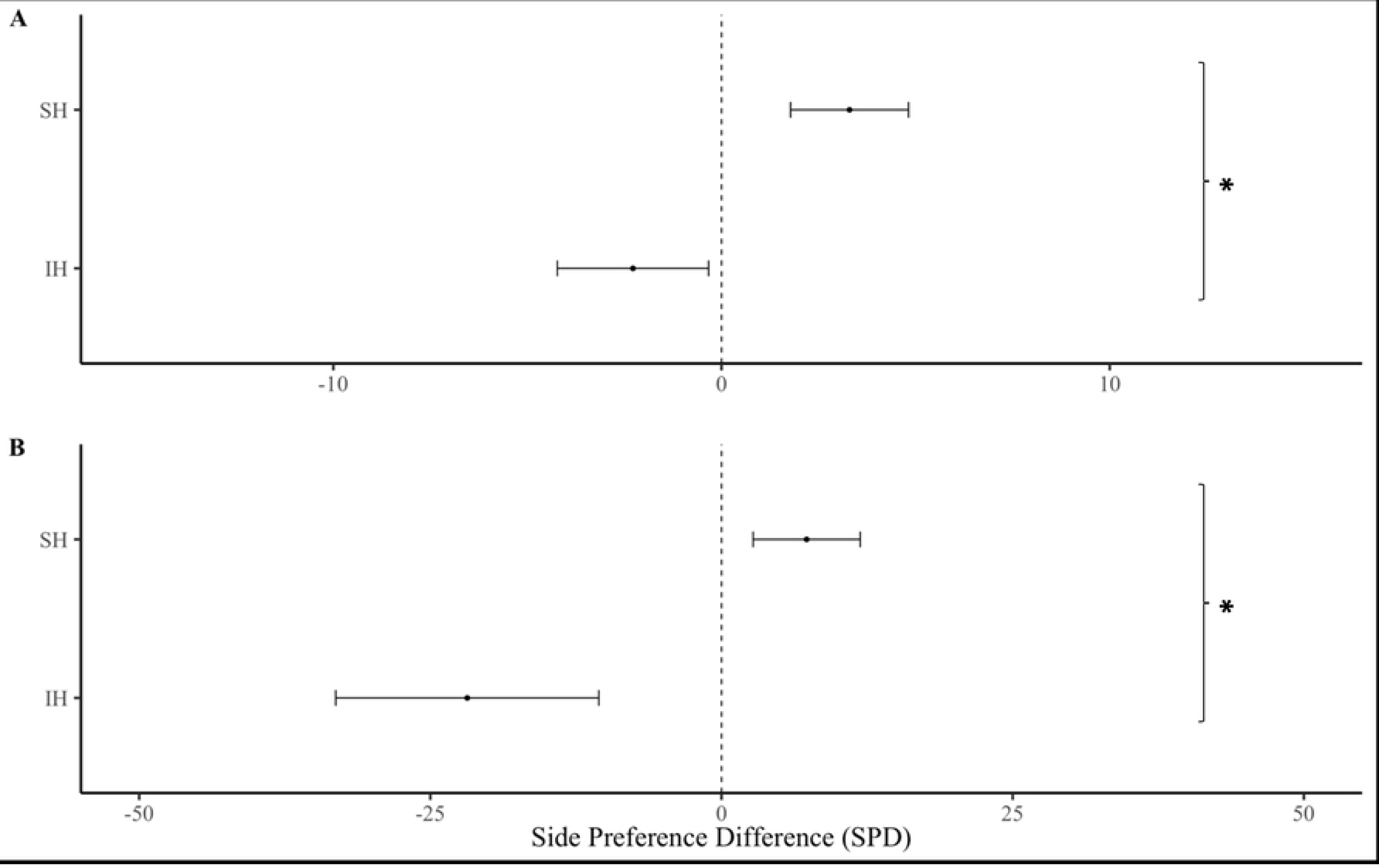
Side preference difference (SPD) of females from different housing treatments. Showing **(A)** over one min and **(B)** over five mins. Error bars show mean ± SE. Asterisks show significant differences.

**Figure 4.**
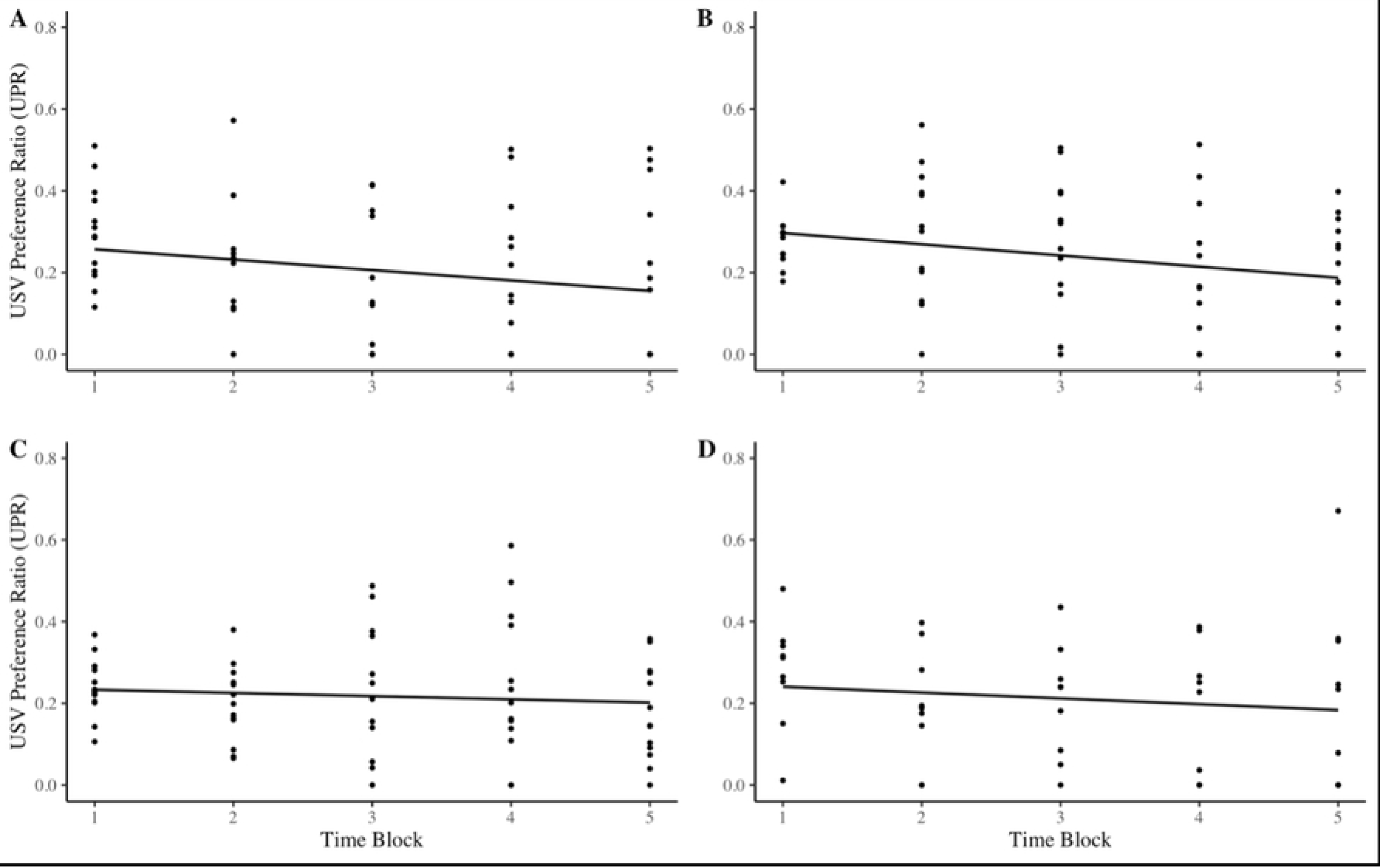
Females’ preferences for male USVs over one-min intervals. Results of females socially housed and reared **(A)** with or **(B)** without a father, and individually housed and reared **(C)** with or **(D)** without father. Preferences measure using USV preference ratio (UPR) and mean linear regression lines shown.

### Sexual receptivity

There was no significant interaction between female sexual receptive stage and their preference for either the USV playback stimulus or the control over five min (F _(1,46)_ = 0.10, p=0.750), however there was a significant interaction over the first min (F _(1,46)_ = 4.29, p=0.044), non-receptive females spent more time in the USV chamber than receptive females. There was no significant difference in SPD from females in different receptive stages over five mins (F _(1,43)_ = 0.66, p=0.422) or one min (F _(1,44)_ = 3.16, p=0.084).

There was no significant interaction between housing treatment and receptive stage on SPD over five min (F _(1,43)_ = 0.10, p=0.753) or one min (F _(1,43)_ = 3.32, p=0.076). Binomial tests revealed a higher proportion of receptive females over non-receptive females within the IH treatment group (p=0.017), though this was not the case for SH females (p= 0.327). To determine whether this difference in ratio was subject to housing conditions, we followed this result up with a binomial regression that revealed these proportions were not significantly different from one another (F _(1,46)_ = 1.35, p=0.250). When exploring the effects of receptive stage on SPD amongst SH, we found that non-receptive females from the SH treatment group had a significantly higher SPD than receptive females (F _(1,23)_ = 8.4, p<0.01), although this pattern was not observed for IH females (F _(1,18)_ = 0.11, p=0.746).

**Figure 5.**
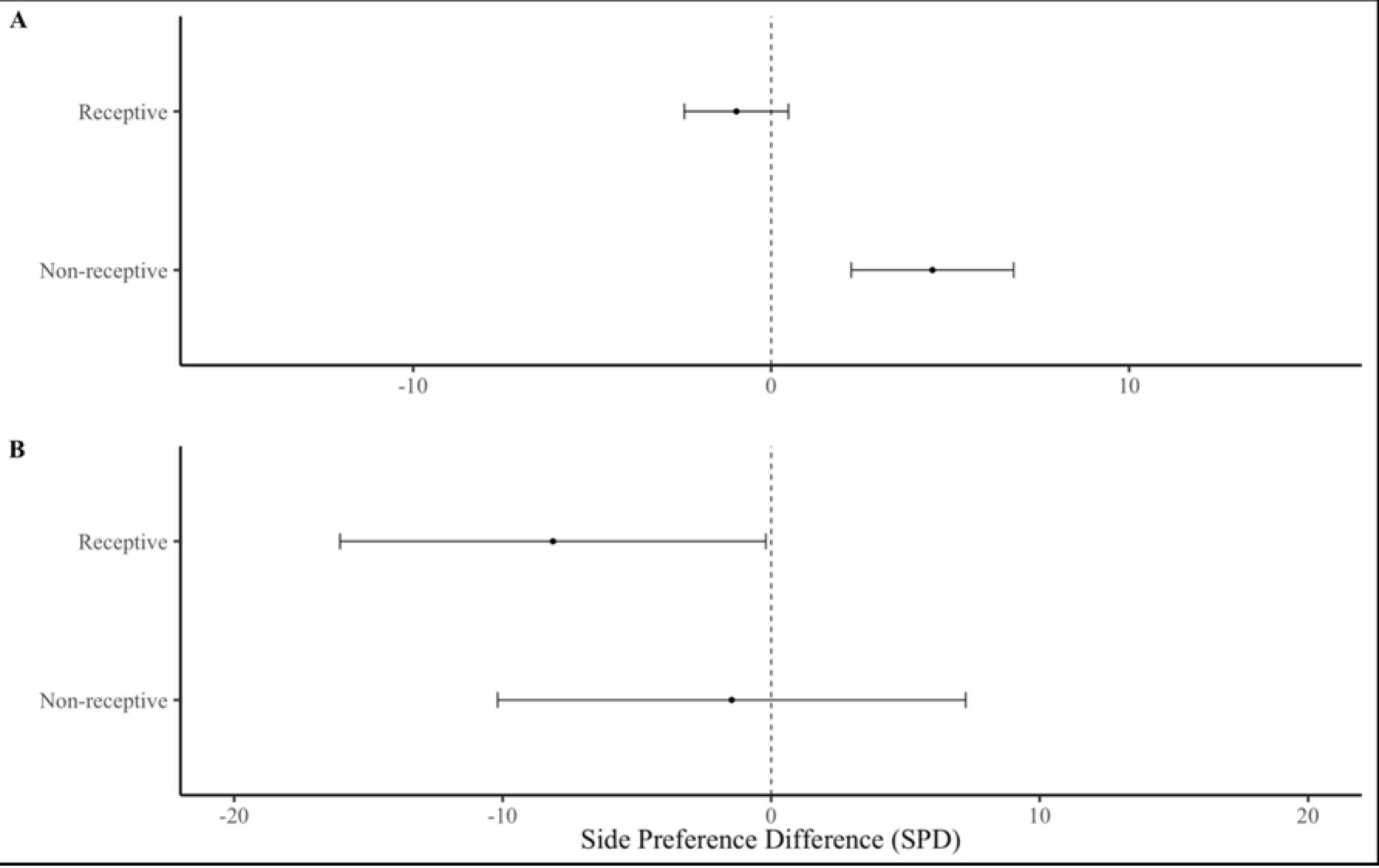
Side preference difference of females in different receptive stages. **(A)** over one min and **(B)** over five mins. Error bars show mean ± SE. Asterisks show significant differences.

**Figure 6.**
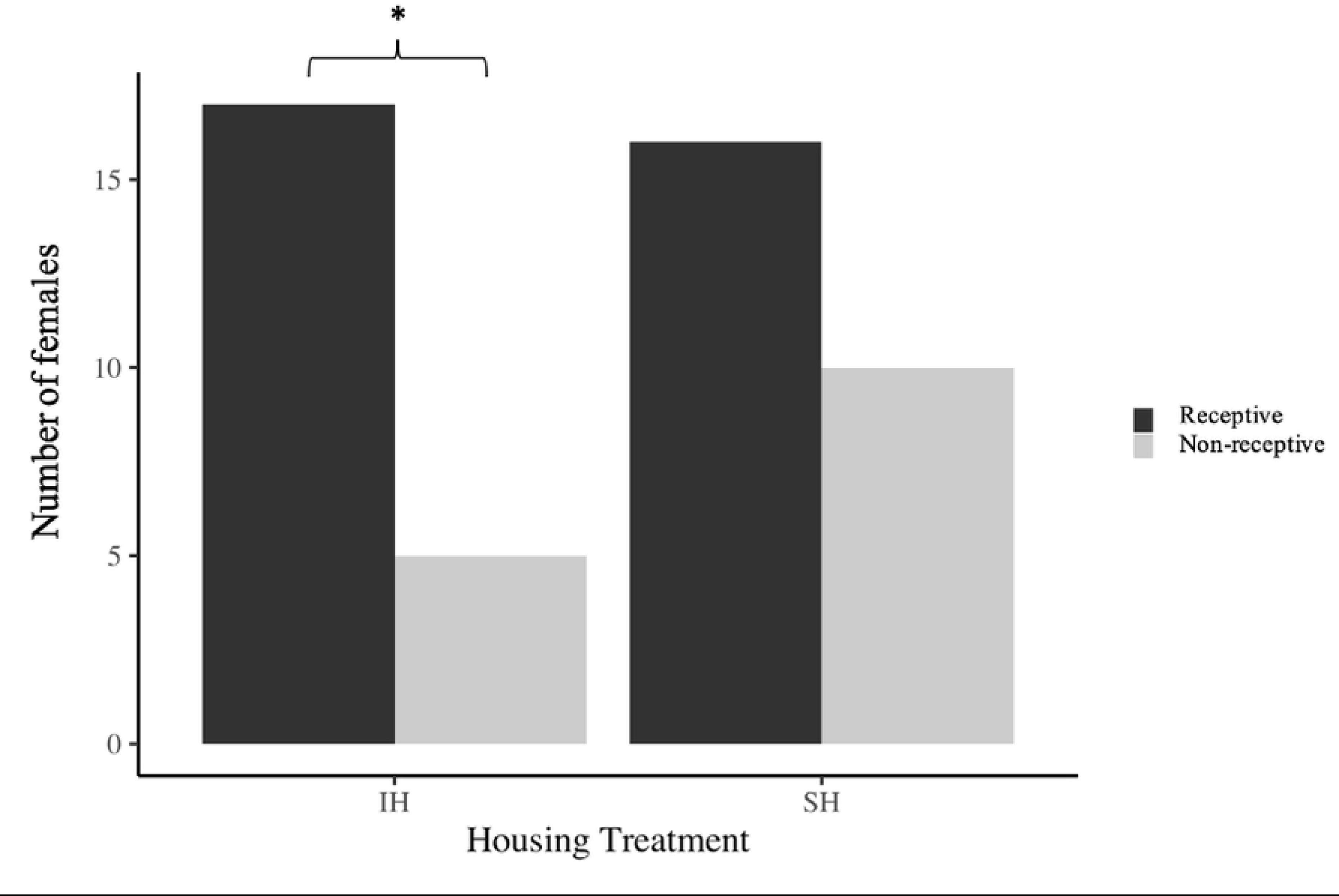
Number of females in sexually receptive and non-receptive stages comparing individually housed (IH) versus socially housed (SH) conditions. Asterisks show significant differences.

**Figure 7.**
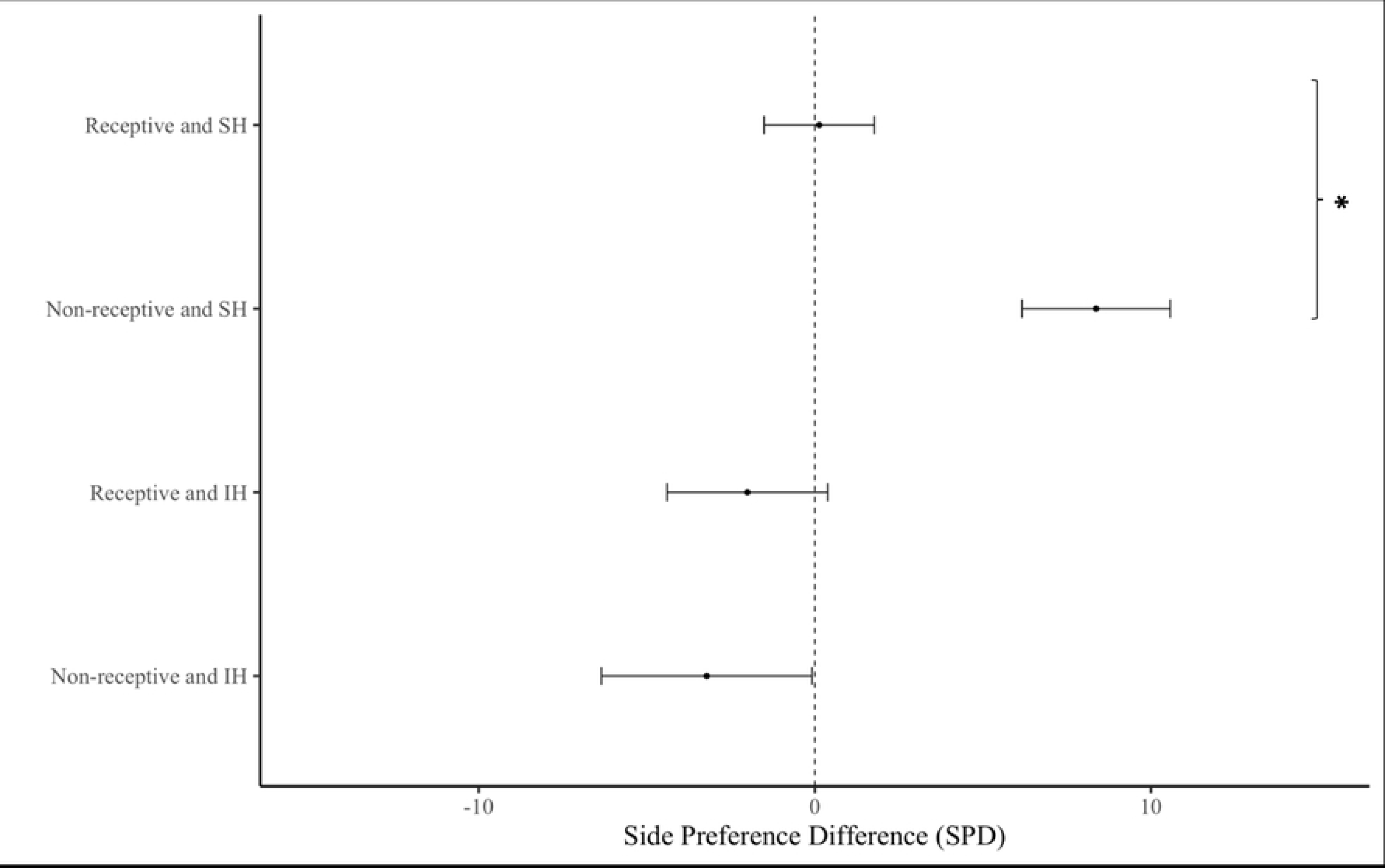
Side preference difference of females in different receptive stages across different housing conditions. Error bars show mean ± SE. Asterisks show significant differences.

## Discussion

Overall, female subjects were not more attracted to playbacks of male USVs than controls, which contrasts with previous studies [4, 24], though we used different methods (e.g., background noise as a control rather than silence or white noise, and no male odor as an enhancing stimulus). Yet, SH females showed preferences for male USVs versus controls, whereas IH females showed the opposite response, which explains why there was no overall effect. We found no evidence that paternal exposure influenced female attraction to male USVs; however, our results do not rule out the possibility that PE enhances female preferences if presented with USVs of an unfamiliar versus familiar male (sexual imprinting hypothesis) [28]. Sexual receptivity (estrous stage) influenced female attraction for male USVs, as females in non-receptive estrous stages were more attracted to male USV playbacks than those in sexually receptive estrous stages [28]. An interaction between housing condition and estrous cycle on USV preferences was not detected, potentially due to insufficient sample size or an unbalanced design, and thus, we cannot rule out that these factors work in combination. Our results indicate that female attraction to male USVs depends upon social housing and estrous stage but does not appear to require paternal exposure. Below we address each of our main findings in more detail.

### Social Housing

Social housing not only influenced female responses in our study, it was necessary for females to show attraction towards playbacks of male USVs. SH females showed preferences for USVs over both the first minute and the entire five minutes of the trial, whereas previous playback studies [4, 26, 28-29] found that females rapidly habituate to playbacks over time. In contrast, IH females showed a preference for background noise over male USVs, responding as if they perceived male vocalizations as aversive. Females from all social backgrounds showed habituation and spent less time in the USV chamber over time. It is unclear why females quickly habituate to USV playbacks, though they may lose interest because they quickly realize that no mice are present.

In contrast, a previous study found social housing had no effect on female attraction to playbacks of USVs [32], and IH females and SH females (housed with females but not females housed with males) showed increased attraction to USVs over time rather than habituation [4, 26, 28-29]. The difference in our results could be explained one or more methodological differences, i.e., we studied 48 wild female *Mus musculus musculus* (versus 18 female CBA mice), whose estrous stages were determined, using recordings from 12 males (versus one individual male) and background noise as a control (versus white noise), and we tested female discrimination in a Y-maze (versus a three-chamber assay), and the rearing and social housing conditions of the mice also differed.

Previous studies with laboratory mice have found that social housing induces several changes that could explain our results by altering: (1) female sexual receptivity (i.e., IH females are less likely to display sexually receptive lordosis posture during sexual interactions than SH females) [36]; (2) auditory perception of USVs (i.e., female IH mice require more time to learn to discriminate a pure tone, though differences disappeared after a few training sessions) [50]; (3) auditory neurons of the brain (females [38, 41], males [51]); and functional neuro-connectivity (i.e., dependencies among remote neurophysiological events [52] was is lower in IH versus SH male mice when exposed to a playback of a female broadband vocalization [40]). Social housing has also been shown to influence hormone levels (i.e., IH females and males have lower corticosterone levels than SH mice [53-54] (but see [55]), and other behaviors, such as hyperlocomotion and increased anxiety/impulsivity in an elevated plus maze test, impaired learning (object recognition) and other aspects of cognition [56], though not always [57]). However, it is premature to generalize these results, as the effects of social housing are often strain-, assay, and sex-specific [58], and can depend upon the timing and duration of the treatments.

The effects of IH on female responses to USVs and other behaviors might be a laboratory artifact due to rearing animals in artificial social conditions. For example, IH results in increasing USV emission due to inducing same-sex mounting during social interactions (male-male mounting in CBA mice [59]; female-female mounting in B6 mice [60]). However, contrary to what is often assumed, not all changes from IH are necessarily pathological nor maladaptive. Female mice in the wild potentially benefit by being cautious when approaching unfamiliar mice when they are solitary, such as during dispersal or in low population densities. Similarly, females can potentially benefit by altering their estrous cycling depending upon the number and sex of conspecifics [61-62] (see below). Little is known about the fitness consequences of changes induced by socio-sexual experience. One study found that keeping male house mice in IH versus SH (with females) had no detectable affect their mating or reproductive success in a female mate choice experiment [63]. Regardless, mice kept in IH versus SH show many unexpected differences in their vocal communication, including USV emission [19, 59-60], auditory perception [50], auditory neural pathways [39-41], and responses to USVs, which researchers should consider in future studies.

### Paternal exposure

We found no evidence that neonatal paternal exposure influenced females’ preferences for male USV, suggesting that PE is not necessary for females to develop USV preferences. A previous study found evidence that female preferences for male USVs are acquired during early development, and that paternal exposure prior to weaning increases female responsiveness to unfamiliar USV playbacks [28] (whereas in birds, females positively imprint on their fathers’ vocalizations [63-65]). Unlike this previous study, however, we did not compare female preferences for the vocalizations of familiar versus unfamiliar males. Therefore, our results suggest that neonatal exposure to male USVs are unnecessary for females to develop preferences for (unfamiliar) USVs; but they do not rule out sexual imprinting [28]. It is possible that PE had no effect in our study because subjects shared a colony room with other mice and were likely exposed to the USVs of other adult males in neighboring cages. This type of indirect exposure may have been sufficient for females to develop a preference for male vocalizations, and thus, we do not rule out the hypothesis that exposure to USVs influences the development of USV preferences. The subjects in our study shared a colony room with other mice and were likely exposed to the USVs of other adult males in neighboring cages, which may have been sufficient to develop a preference for male vocalizations. Prior to weaning, females in our study were also exposed to their litter mates’ USVs, though pup vocalizations differ from adult males [66], which may be sufficient to induce attraction to male USVs.

### Sexual receptivity

We found that female attraction to male USVs was affected by their sexual receptivity, but only in the first minute and not over the entirety of the trial. We found that non-receptive females (diestrus and metestrus) spent a significantly longer time in the USV chamber than receptive females (proestrus and estrous). Our result is consistent with a study on laboratory mice that found only diestrus females showed preferences for male USVs [28, 67]. Females might be attracted to male USVs before estrus because they begin assessing potential mates during this period (wild house mice usually take many days and often many weeks to mate, especially in seminatural conditions [47-48]). However, this interpretation does not explain why females are not attracted to male USVs during estrus. It has been shown that social investigation towards unfamiliar males is enhanced by moderate levels of estrogen [68] and sexually receptive (estrus) females in other species show stronger preferences for male odor than non-receptive females [69-71], but why female responses for male USV are reversed is unclear. Asaba et al [28] proposed that females have two systems for mate preferences: the first is a preference for masculine males (social rank or higher testosterone) in the estrus phase [71-73]; the second is inbreeding avoidance in the diestrus stage [28, 67]. A previous study found that females in estrus were preferentially attracted to the USVs of unfamiliar non-kin over those of familiar kin [4], though they were not compared with anestrus females and this result is based on a smaller sample size (n=10) to the previously mentioned study [28] (n=40). Whilst in our study, the number of vocalizations (which may be influenced by testosterone [74]) was standardized, all recordings came from unrelated males. Thus, to test the inbreeding avoidance hypothesis, future studies comparing female USV preference to familiar vs unfamiliar kin amongst females of different estrous stages is required. Moreover, studies are needed to manipulate estrous stage experimentally to test its effects on female responses to male USVs.

### Social housing and sexual receptivity

In addition to testing the effects of housing, paternal exposure, and estrous cycle on female proclivity to investigate male USVs, we also investigated whether these factors interact. We found that a higher proportion of IH females were in receptive estrous stages than non-receptive estrous stages, although this was not the case for SH females. Amongst SH females, those in non-receptive estrous stages showed greater USV preferences than those in receptive stages, but there was no effect of sexual receptivity on USV preference amongst IH females. There was no significant interaction between housing type and estrous stage on female USV preference when investigating the entire sample population.

The influence of housing conditions on estrous cycling has been well documented in house mice, as females housed in groups and isolated from males show suppression or prolongation of estrous cycles (Lee-Boot effect) [61]. Isolated females tend to display short estrous cycles of 4–6 days [75], whereas pair-housed females are lengthened slightly, and group-housed females (4-6 individuals) have even longer cycles [76] with complete suppression occurring once all-female groups become even larger [77]. To our knowledge, no study has previously examined whether the Lee-Boot effect influences female preference for male USVs. Exposure to male scent induces estrous cycling in wild house mice [62], and we exposed females to male bedding 48 h before trials to ensure cycling. Therefore, our result suggests that the estrous cycles of IH females were shorter than those of SH females, consistent with the Lee-Boot effect. However, further investigation of the regression analysis suggests that these perceived differences in estrus ratio are not due to housing treatment and so we are unable to make strong conclusions.

Although the interaction between social housing and estrous cycle was not statistically significant (p=0.076), our results suggest that housing condition and estrous stage work in combination, rather than independently to affect female USV preferences. It will be difficult to disentangle their independent effects, and the Lee-Boot effect has been shown in wild house mice [62], as well as laboratory mice [61, 75-77]. Many of the differences in socio-sexual behaviors found between SH and IH females in other studies [36, 38, 50] did not report the estrous stage of the mice [36, 50] or an unbalanced design was used [38]. In our study, we used 26 SH (16 receptive, 10 non-receptive) and 22 IH females (17 receptive, 5 receptive), and thus studies using a larger sample size and a balanced design – as well as experimentally manipulating estrous stage – are needed to investigate this potential interaction.

### Conclusions and Future Directions

Our results provide the first evidence to our knowledge that female house mice are more likely to show attraction towards male USVs if they are socially housed, whereas individually housed females are more likely to avoid them. We found no effect of neonatal paternal exposure on female preference for male USV playbacks, though this result does not rule out sexual imprinting (and may be due to females in our study being exposed to auditory cues of adult males in our colony room). We found that non-receptive females show greater preferences for male playbacks than sexually receptive females, as previously reported [28], and this result was most pronounced amongst SH females and did not appear amongst IH females.

Our findings contribute to a growing number of studies showing that social housing [33] and estrous stage [46] influence the behavior of house mice, and raise the possibility that these factors may interact. Studies are now needed to explore the ecological relevance of these results, such as by varying the number individuals, density, and sex ratio in natural or seminatural conditions [47], and not merely comparing the effects of IH versus SH in small cages [45]. Future research is also needed to investigate the effects of the timing and duration of social housing and estrous stage on behavior (i.e., ontogeny), and underlying proximate mechanisms, including endocrine responses [53], sensory perception [37], neural pathways [39-41] and their adaptive consequences, such as altering sexual motivation [36] and mating success and reproductive success [63].

Although results from studies on social housing have prompted recommendations to stop the use of IH in animal research [78], such prescriptions are too general and overly simplistic [79], especially for wild house mice, which are highly territorial and aggressive towards male cage mates. Furthermore, the various effects of IH are not necessarily pathological, contrary to what is generally assumed (e.g., solitary females may benefit by being cautious about approaching the USVs of unfamiliar mice). To better understand why many behaviors depend upon social housing and experience, more studies are needed to determine the fitness consequences of altering behavior in response to social experiences and social information (e.g., social competence) [80].

Finally, publications of studies on rodents should provide more specifics about housing conditions and estrous status, as our results show that these sources of variation can influence behavior. It is important to note that if social housing had not been recorded in our study, we would have erroneously concluded that females show no attraction to male USVs, i.e., a false negative. Thus, improving the reporting of variables such as these that have been shown to influence behavior and physiology should help to address the replication crisis [42-45].

## Supporting Information

**S1 File (PDF) Protocol for conducting playback experiments**

**S2 File (txt) Data file: USV Preferences Raw Data-One Min Analysis**

**S3 File (txt) Data file: USV Preferences Raw Data-Five Min Analysis**

**S4 File (txt) Data file: USV Preferences Raw Data-Habituation**

## Acknowledgments

We are grateful to Simon Wölfl and Laura Liebhart for help with data collection and Adelheid Sasse with animal care.

## Authors’ Contribution

**Funding acquisition:** Dustin Penn, Sarah Zala

**Conceptualization:** Dustin Penn, Sarah Zala

**Project administration:** Sarah Zala

**Formal analyses:** Jakob Beck

**Investigation:** Bettina Wernisch, Teresa Klaus

**Methodology:** Sarah Zala, Bettina Wernisch, Teresa Klaus, Dustin Penn

**Writing:** Jakob Beck (original draft), Dustin Penn, Sarah Zala

**Data curation:** Bettina Wernisch

## References

1. Nicolakis D, Marconi MA, Zala SM, Penn DJ. Ultrasonic vocalizations in house mice depend upon genetic relatedness of mating partners and correlate with subsequent reproductive success. Front Zool. 2020;17: 1–9.

2. Asaba A, Osakada T, Touhara K, Kato M, Mogi K, Kikusui T. Male mice ultrasonic vocalizations enhance female sexual approach and hypothalamic kisspeptin neuron activity. Horm Behav. 2017;94: 53–60.

3. Hoffmann F, Musolf K, Penn DJ. Ultrasonic courtship vocalizations in wild house mice: spectrographic analyses. J Ethol. 2012;30: 173–180.

4. Musolf K, Hoffmann F, Penn DJ. Ultrasonic courtship vocalizations in wild house mice, Mus musculus musculus. Anim Behav. 2010;79(3): 757–764.

5. Portfors CV, Perkel DJ. The role of ultrasonic vocalizations in mouse communication. Curr Opin Neurobiol. 2014;28: 115-20.

6. Heckman J, McGuinness B, Celikel T, Englitz B. Determinants of the mouse ultrasonic vocal structure and repertoire. Neurosci Biobehav Rev. 2016;65: 313–325.

7. Nyby J. Ultrasonic vocalizations during sex behavior of male house mice (Mus musculus): a description. Behav Neural Biol. 1983;39(1): 128–134.

8. Matsumoto YK, Okanoya K. Phase-specific vocalizations of male mice at the initial encounter during the courtship sequence. PloS one. 2016;11(2): 0147102.

9. Holy TE, Guo Z. Ultrasonic songs of male mice. PLoS Biol. 2005;3(12): 386.

10. Zala SM, Reitschmidt D, Noll A, Balazs P, Penn DJ. Automatic mouse ultrasound detector (A-MUD): A new tool for processing rodent vocalizations. PloS one. 2017;12(7): 0181200.

11. Abbasi R, Balazs P, Marconi MA, Nicolakis D, Zala SM, Penn DJ. Capturing the songs of mice with an improved detection and classification method for ultrasonic vocalizations (BootSnap). PLoS Comput Biol. 2022;18(5): 1010049.

12. Neunuebel JP, Taylor AL, Arthur BJ, Egnor SR. Female mice ultrasonically interact with males during courtship displays. eLife. 2015;4: 06203.

13. Zala SM, Reitschmidt D, Noll A, Balazs P, Penn DJ. Sex-dependent modulation of ultrasonic vocalizations in house mice (*Mus musculus musculus*). PloS one. 2017;12(12): 0188647.

14. Ivanenko A, Watkins P, van Gerven MAJ, Hammerschmidt K, Englitz B. Classifying sex and strain from mouse ultrasonic vocalizations using deep learning. PLoS Comput Biol. 2020;16(6): 1007918.

15. Hoffmann F, Musolf K, Penn DJ. Spectrographic analyses reveal signals of individuality and kinship in the ultrasonic courtship vocalizations of wild house mice. Physiol Behav. 2012;105(3): 766–771.

16. Marconi MA, Nicolakis D, Abbasi R, Penn DJ, Zala SM. Ultrasonic courtship vocalizations of male house mice contain distinct individual signatures. Anim Behav. 2020;169: 169–197.

17. Melotti L, Siestrup S, Peng M, Vitali V, Dowling D, von Kortzfleisch VT, Bračić M, Sachser N, Kaiser S, Richter SH. Individuality, as well as genetic background, affects syntactical features of courtship songs in male mice. Anim Behav. 202;180: 179–196.

18. Lopes PC, König B. Choosing a healthy mate: sexually attractive traits as reliable indicators of current disease status in house mice. Anim Behav. 2016;111: 119–126.

19. Chabout J, Serreau P, Ey E, Bellier L, Aubin T, Bourgeron T, Granon S. Adult male mice emit context-specific ultrasonic vocalizations that are modulated by prior isolation or group rearing environment. PloS one. 2012;7(1): 29401.

20. Zala SM, Nicolakis D, Marconi MA, Noll A, Ruf T, Balazs P, Penn DJ. Primed to vocalize: Wild-derived male house mice increase vocalization rate and diversity after a previous encounter with a female. PloS one. 2020;15(12): 0242959.

21. Burke K, Screven LA, Dent ML. CBA/CaJ mouse ultrasonic vocalizations depend on prior social experience. PLoS One. 2018;13(6): 0197774.

22. Kanno K, Kikusui T. Effect of sociosexual experience and aging on number of courtship ultrasonic vocalizations in male mice. Zoolog Sci. 2018;35(3): 208–214.

23. Hanson JL, Hurley LM. Female presence and estrous state influence mouse ultrasonic courtship vocalizations. PloS one. 2012;7(7): 40782.

24. Hammerschmidt K, Radyushkin K, Ehrenreich H, Fischer J. Female mice respond to male ultrasonic ‘songs’ with approach behaviour. Biol Lett. 2009;5(5): 589–592.

25. Chabout J, Sarkar A, Dunson DB, Jarvis ED. Male mice song syntax depends on social contexts and influences female preferences. Front Behav Neurosci. 2015;9: 76.

26. Hammerschmidt K, Radyushkin K, Ehrenreich H, Fischer J. The structure and usage of female and male mouse ultrasonic vocalizations reveal only minor differences. PloS one. 2012;7(7): 41133.

27. Musolf K, Meindl S, Larsen AL, Kalcounis-Rueppell MC, Penn DJ. Ultrasonic vocalizations of male mice differ among species and females show assortative preferences for male calls. PLoS one. 2015;10(8): 0134123.

28. Asaba A, Okabe S, Nagasawa M, Kato M, Koshida N, Osakada T, Mogi K, Kikusui T. Developmental social environment imprints female preference for male song in mice. PloS one. 2014;9(2): 87186.

29. Shepard KN, Liu RC. Experience restores innate female preference for male ultrasonic vocalizations. Genes Brain Behav. 2011;10(1): 28–34.

30. Hurst JL. Female recognition and assessment of males through scent. Behav Brain Res. 2009;200(2): 295–303.

31. Brown RE. Effects of social isolation in adulthood on odor preferences and urine-marking in male rats. Behav Neural Biol. 1985;44(1): 139–143.

32. Screven LA, Dent ML. Preference in female laboratory mice is influenced by social experience. Behav Processes. 2018;157: 171–179.

33. Arakawa H. Ethological approach to social isolation effects in behavioral studies of laboratory rodents. Behav Brain Res. 2018;341: 98–108.

34. Liu ZW, Yu Y, Lu C, Jiang N, Wang XP, Xiao SY, Liu XM. Postweaning isolation rearing alters the adult social, sexual preference and mating behaviors of male CD-1 mice. Front Behav Neurosci. 2019;13: 21.

35. Amistislavskaya TG, Bulygina VV, Tikhonova MA, Maslova LN. Social isolation during peri-adolescence or adulthood: effects on sexual motivation, testosterone, and corticosterone response under conditions of sexual arousal in male rats. Chin J Physiol. 2013;56(1): 36–43.

36. Kercmar J, Tobet SA, Majdic G. Social isolation during puberty affects female sexual behavior in mice. Front Behav Neurosci. 2014;8: 337.

37. Keesom SM, Hurley LM. Silence, solitude, and serotonin: neural mechanisms linking hearing loss and social isolation. Brain Sci. 2020;10(6): 367.

38. Davis SE, Sansone JM, Hurley LM. Postweaning isolation alters the responses of auditory neurons to serotonergic modulation. Integr Comp Biol. 2021;61(1): 302–315.

39. Dixon AK. The social behaviour of mice and its sensory control. In: Hedrich HJ, Bullock G, editors. The laboratory mouse. Oxford: Academic Press; 2004. pp. 287-298.

40. Petersen CL, Davis SE, Patel B, Hurley LM. Social experience interacts with serotonin to affect functional connectivity in the social behavior network following playback of social vocalizations in mice. eNeuro. 2021;8(2).

41. Keesom SM, Morningstar MD, Sandlain R, Wise BM, Hurley LM. Social isolation reduces serotonergic fiber density in the inferior colliculus of female, but not male, mice. Brain Res. 2018;1694: 94–103.

42. von Kortzfleisch VT, Ambrée O, Karp NA, Meyer N, Novak J, Palme R, Rosso M, Touma C, Würbel H, Kaiser S, Sachser N. Do multiple experimenters improve the reproducibility of animal studies?. PLoS Biol. 2022;20(5): 3001564.

43. Voelkl B, Altman NS, Forsman A, Forstmeier W, Gurevitch J, Jaric I, Karp NA, Kas MJ, Schielzeth H, Van de Casteele T, Würbel H. Reproducibility of animal research in light of biological variation. Nat Rev Neurosci. 2020;21(7): 384–393.

44. Crabbe JC, Wahlsten D, Dudek BC. Genetics of mouse behavior: interactions with laboratory environment. Science. 1999;284(5420): 1670-1672.

45. Saré RM, Lemons A, Smith CB. Behavior testing in rodents: highlighting potential confounds affecting variability and reproducibility. Brain Sci. 2021;11(4): 522.

46. Chari T, Griswold S, Andrews NA, Fagiolini M. The stage of the estrus cycle is critical for interpretation of female mouse social interaction behavior. Front Behav Neurosci. 2020;14: 113.

47. Luzynski KC, Nicolakis D, Marconi MA, Zala SM, Kwak J, Penn DJ. Pheromones that correlate with reproductive success in competitive conditions. Sci Rep. 2021;11(1): 21970.

48. Thoß M, Luzynski KC, Enk VM, Razzazi-Fazeli E, Kwak J, Ortner I, Penn DJ. Regulation of volatile and non-volatile pheromone attractants depends upon male social status. Sci Rep. 2019;9(1): 489.

49. R Core Team. R: A Language and Environment for Statistical Computing. Vienna, Austria; 2016. Available from: https://www.R-project.org/.

50. Screven LA, Dent ML. Perception of ultrasonic vocalizations by socially housed and isolated mice. eNeuro. 2019;6(5).

51. Keesom SM, Sloss BG, Erbowor-Becksen Z, Hurley LM. Social experience alters socially induced serotonergic fluctuations in the inferior colliculus. J Neurophysiol. 2017;118(6): 3230–3241.

52. Friston KJ. Functional and effective connectivity: a review. Brain Connect. 2011;1(1): 13–36.

53. Kamakura R, Kovalainen M, Leppäluoto J, Herzig KH, Mäkelä KA. The effects of group and single housing and automated animal monitoring on urinary corticosterone levels in male C57 BL/6 mice. Physiol Rep. 2016;4(3): 12703.

54. Martin AL, Brown RE. The lonely mouse: verification of a separation-induced model of depression in female mice. Behav Brain Res. 2010;207(1): 196–207.

55. Hunt C, Hambly C. Faecal corticosterone concentrations indicate that separately housed male mice are not more stressed than group housed males. Physiol Behav. 2006;87(3): 519–526.

56. Koike H, Ibi D, Mizoguchi H, Nagai T, Nitta A, Takuma K, Nabeshima T, Yoneda Y, Yamada K. Behavioral abnormality and pharmacologic response in social isolation-reared mice. Behav Brain Res. 2009;202(1): 114–121.

57. Arndt SS, Laarakker MC, van Lith HA, van der Staay FJ, Gieling E, Salomons AR, van’t Klooster J, Ohl F. Individual housing of mice—impact on behaviour and stress responses. Physiol Behav. 2009;97(3-4):385–393.

58. Voikar V, Polus A, Vasar E, Rauvala H. Long-term individual housing in C57BL/6J and DBA/2 mice: assessment of behavioral consequences. Genes Brain Behav. 2005;4(4): 240–252.

59. Keesom SM, Finton CJ, Sell GL, Hurley LM. Early-life social isolation influences mouse ultrasonic vocalizations during male-male social encounters. PloS one. 2017;12(1): 0169705.

60. Zhao X, Ziobro P, Pranic NM, Chu S, Rabinovich S, Chan W, Zhao J, Kornbrek C, He Z, Tschida KA. Sex-and context-dependent effects of acute isolation on vocal and non-vocal social behaviors in mice. PloS one. 2021;16(9): 0255640.

61. van der Lee S, Boot L. Spontaneous pseudopregnancy in mice. II. Acta Physiol Pharmacol Neerl. 1956;5(2): 213–215.

62. Wölfl S, Zala SM, Penn DJ. Male scent but not courtship vocalizations induce estrus in wild female house mice. Physiol Behav. 2023;259: 114053.

63. Thonhauser KE, Raffetzeder A, Penn DJ. Sexual experience has no effect on male mating or reproductive success in house mice. Scientific Reports. 2019;9(1): 1–11.

64. Clayton NS. Song discrimination learning in zebra finches. Anim Behav. 1988;36(4): 1016–1024.

65. Miller DB. Long-term recognition of father’s song by female zebra finches. Nature. 1979;280(5721): 389–391.

66. Grimsley JM, Monaghan JJ, Wenstrup JJ. Development of social vocalizations in mice. PloS one. 2011;6(3): 17460.

67. Asaba A, Kato M, Koshida N, Kikusui T. Determining ultrasonic vocalization preferences in mice using a two-choice playback test. J Vis Exp. 2015;(103): 53074.

68. Yano S, Sakamoto KQ, Habara Y. Estrus cycle-related preference of BALB/c female mice for C57BL/6 males is induced by estrogen. J Vet Med Sci. 2012;74(10): 1311– 1314.

69. Doty RL. Odor preferences of female Peromyscus maniculatus bairdi for male mouse odors of P. m. bairdi and P. leucopus noveboracensis as a function of estrous state. J Comp Physiol Psychol. 1972;81(2): 191–197.

70. Johnston RE. Olfactory preferences, scent marking, and “proceptivity” in female hamsters. Horm Behav. 1979;13(1): 21–39.

71. Pasch B, George AS, Campbell P, Phelps SM. Androgen-dependent male vocal performance influences female preference in Neotropical singing mice. Anim Behav. 2011;82(2): 177–83.

72. Baum MJ, Keverne EB. Sex difference in attraction thresholds for volatile odors from male and estrous female mouse urine. Horm Behav. 2002;41(2): 213–219.

73. Xiao K, Kondo Y, Sakuma Y. Sex-specific effects of gonadal steroids on conspecific odor preference in the rat. Hormones and Behavior. 2004;46(3): 356–361.

74. Nunez AA, Nyby J, Whitney G. The effects of testosterone, estradiol, and dihydrotestosterone on male mouse (Mus musculus) ultrasonic vocalizations. Horm Behav. 1978;11(3): 264–272.

75. Ma W, Miao Z, Novotny MV. Role of the adrenal gland and adrenal-mediated chemosignals in suppression of estrus in the house mouse: the lee-boot effect revisited. Biol Reprod. 1998;59(6): 1317–1320.

76. Champlin AK. Suppression of oestrus in grouped mice: the effects of various densities and the possible nature of the stimulus. Reprod. 1971;27(2): 233–241.

77. Whitten WK. Occurrence of anoestrus in mice caged in groups. J Endocrinol. 1959;18(1): 102–107.

78. Lewejohann L, Schwabe K, Häger C, Jirkof P. Impulse for animal welfare outside the experiment. Lab Anim. 2020;54(2): 150–158.

79. Kappel S, Hawkins P, Mendl MT. To group or not to group? Good practice for housing male laboratory mice. Animals. 2017;7(12): 88.

80. Taborsky B, Oliveira RF. Social competence: an evolutionary approach. Trends Ecol Evol. 2012;27(12): 679–688.

